# MeLSI: Metric Learning for Statistical Inference in Microbiome Community Composition Analysis

**DOI:** 10.64898/2025.12.04.692328

**Authors:** Nathan Bresette, Aaron C. Ericsson, Carter Woods, Ai-Ling Lin

## Abstract

Microbiome beta diversity analysis relies on distance-based methods including PERMANOVA combined with fixed ecological distance metrics (Bray-Curtis, Euclidean, Jaccard, and UniFrac), which treat all microbial taxa uniformly regardless of their biological relevance to community differences. This “one-size-fits-all” approach may miss subtle but biologically meaningful patterns in complex microbiome data. We present MeLSI (Metric Learning for Statistical Inference), a novel machine learning framework that learns data-adaptive distance metrics optimized for detecting community composition differences in multivariate microbiome analyses. MeLSI employs an ensemble of weak learners using bootstrap sampling, feature subsampling, and gradient-based optimization to learn optimal feature weights, combined with rigorous permutation testing for statistical inference. The learned metrics can be used with PERMANOVA for hypothesis testing and with Principal Coordinates Analysis (PCoA) for ordination visualization. Comprehensive validation on synthetic benchmarks and real datasets shows that MeLSI maintains proper Type I error control while delivering competitive or superior F-statistics when signal structure aligns with CLR-based weighting and, crucially, supplies interpretable feature-weight profiles that clarify which taxa drive group separation. On the Atlas1006 dataset, MeLSI achieved stronger effect sizes than the best traditional methods, and even when performance was comparable, the learned feature weights provided biological insight that fixed metrics cannot supply. MeLSI therefore offers a statistically rigorous tool that augments beta diversity analysis with transparent, data-driven interpretability.

**IMPORTANCE:** Understanding which microbes differ between groups of interest could reveal therapeutic targets and diagnostic biomarkers. However, current analysis methods treat all microbes equally (similar to using the same ruler to measure everything, regardless of what matters most). This means subtle but clinically important differences may go undetected, especially when only a few key species drive disease while hundreds of “bystander” species add noise. MeLSI solves this by learning which microbes matter most for each specific comparison. In comparing male and female gut microbiomes, MeLSI identified specific bacterial families driving the differences, providing actionable biological insights that standard methods miss. This capability is particularly crucial for detecting early disease biomarkers, where differences are subtle and masked by biological variability. By telling researchers not just whether groups differ, but which specific microbes drive those differences, MeLSI accelerates the path from microbiome data to testable biological hypotheses and clinical applications.

## INTRODUCTION

### The microbiome and human health

The human microbiome, the complex community of microorganisms inhabiting our bodies, plays fundamental roles in health and disease (1, 2). Recent advances in high-throughput sequencing technologies have enabled comprehensive profiling of microbial communities, revealing associations between microbiome composition and diverse conditions including inflammatory bowel disease, obesity, diabetes, and neurological disorders (3, 4). A central question in microbiome research is comparing overall microbial community composition between groups of interest, typically assessed through beta diversity analysis, which studies compositional differences between samples.

### Current approaches and their limitations

Microbiome beta diversity analysis predominantly relies on distance-based multivariate methods including PERMANOVA (Permutational Multivariate Analysis of Variance) combined with fixed ecological distance metrics (5, 6). Commonly used metrics include Bray-Curtis dissimilarity, Euclidean distance, Jaccard index, and phylogenetically-informed metrics including UniFrac (7). These approaches have proven valuable for hypothesis testing about community differences and visualization through ordination methods such as Principal Coordinates Analysis (PCoA) (8).

However, fixed distance metrics suffer from a fundamental limitation. They apply the same mathematical formula to all datasets, treating all microbial taxa with equal importance regardless of their biological relevance to the specific research question (9). For instance, Bray-Curtis dissimilarity equally weights all taxa based on their relative abundances, while Euclidean distance treats all features identically. This “one-size-fits-all” approach may fail to capture subtle but biologically meaningful differences when only a subset of taxa drive group separation (10).

Furthermore, microbiome data presents unique analytical challenges including high dimensionality (often hundreds to thousands of taxa), compositionality (relative abundances sum to a constant), sparsity (many zero counts), and heterogeneous biological signal across features (11). Fixed metrics cannot adapt to these complexities in a data-driven manner.

### The need for statistical rigor

A critical requirement for any beta diversity method is proper statistical inference with controlled Type I error rates (false positive rates). While machine learning approaches often prioritize predictive accuracy, hypothesis testing for community composition differences requires rigorous F-statistic and p-value calculation under the null hypothesis of no group differences (12). Permutation testing provides a non-parametric framework for valid inference that makes minimal distributional assumptions (13), making it particularly suitable for complex microbiome data and distance-based analyses such as PERMANOVA.

### Metric learning: an emerging paradigm

Metric learning, a branch of machine learning, offers a principled approach to address these limitations (14, 15). Rather than using fixed distance formulas, metric learning algorithms learn optimal distance metrics from data by identifying which features contribute most to separating groups of interest. In the context of supervised learning, metric learning algorithms optimize distance functions to maximize between-group distances while minimizing within-group distances (16, 17).

Mahalanobis distance learning (18) learns a positive semi-definite matrix **M** that defines distances as 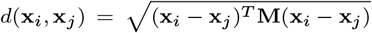. When **M** is diagonal, this reduces to learning feature-specific weights, providing interpretable importance scores (17).

Despite its promise, metric learning has seen limited application in microbiome beta diversity analysis. Previous work has explored metric learning for clinical prediction tasks (19), but not specifically for statistical inference in community composition analysis where rigorous Type I error control is essential.

### Study objectives

We developed MeLSI (Metric Learning for Statistical Inference) to bridge the gap between adaptive machine learning approaches and rigorous statistical inference for microbiome beta diversity and community composition analysis. Our specific objectives were to (1) design an ensemble metric learning framework that learns data-adaptive distance metrics for PERMANOVA and ordination while preventing overfitting, (2) integrate metric learning with permutation testing to ensure valid statistical inference, (3) comprehensively validate Type I error control, statistical power, scalability, parameter sensitivity, and computational efficiency, demonstrate practical utility on real microbiome datasets, and (5) provide interpretable feature importance scores to identify biologically relevant taxa driving community separation.

This paper presents the MeLSI framework, comprehensive validation results, and discussion of its implications for microbiome beta diversity research.

## MATERIALS AND METHODS

### Overview of the MeLSI framework

MeLSI integrates metric learning with permutation-based statistical inference through two main phases:

#### Phase 1: Metric Learning

1. Apply conservative pre-filtering to focus on high-variance features
2. For each of B weak learners:
  - Bootstrap sample the data
  - Subsample features
  - Optimize metric matrix M via gradient descent
3. Combine weak learners via performance-weighted ensemble averaging
4. Compute robust distance matrix using eigenvalue decomposition

#### Phase 2: Statistical Inference

5. Calculate observed F-statistic using the learned metric
6. Generate null distribution via permutation testing (relearn metric on each permutation)
7. Compute permutation-based p-value

Each component addresses specific challenges in microbiome data analysis while maintaining statistical validity. The following sections formalize the mathematical framework and detail each algorithmic component, organized by phase.

### Phase 1: Metric Learning

#### Problem formulation

Let **X** ∈ ℝ^*n*×*p*^ denote a feature abundance matrix with *n* samples and *p* taxa (features), and let **y** = (*y*_1_, …, *y*_*n*_) denote group labels. Our goal is to learn a distance metric optimized for separating groups defined by **y** while ensuring valid statistical inference.

We parameterize the distance metric using a diagonal positive semi-definite matrix **M** ∈ ℝ^*p*×*p*^, where *M*_*jj*_ represents the weight (importance) of feature *j*. The learned Mahalanobis distance between samples *i* and *k* is:

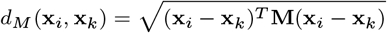

For diagonal M, this simplifies to a weighted Euclidean distance:

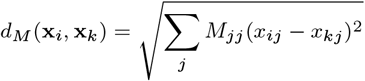

#### Conservative pre-filtering

To improve computational efficiency and reduce noise, MeLSI applies conservative variance-based pre-filtering. For pairwise comparisons, we calculate a feature importance score combining mean differences and variance:

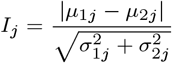

where *µ*_1*j*_ and *µ*_2*j*_ are the mean abundances of feature *j* in groups 1 and 2, and 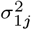 and 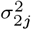 are their variances. We retain the top 70% of features by this importance score, maintaining high statistical power while reducing dimensionality.

For multi-group comparisons (3 or more groups), we use ANOVA F-statistics to rank features and apply the same 70% retention threshold. Critically, this pre-filtering is applied consistently to both observed and permuted data during null distribution generation to avoid bias.

#### Ensemble learning with weak learners

MeLSI constructs an ensemble of *B* weak learners (default *B* = 30) to improve robustness and prevent overfitting. For each weak learner *b*:

1. **Bootstrap sampling:** Draw *n* samples with replacement from the original data to create a bootstrap dataset (**X**_*b*_, **y**_*b*_)
2. **Feature subsampling:** Randomly select *m* = ⌊*p* × *m*_*frac*_⌋ features (default *m*_*frac*_ = 0.8) without replacement
3. **Metric optimization:** Learn **M**_*b*_ on the bootstrapped, subsampled data

The combination of bootstrap sampling (sample-level randomness) and feature subsampling (feature-level randomness) ensures diversity among weak learners, reducing overfitting risk (20).

#### Optimization objective

For each weak learner, we optimize M to maximize between-group distances while minimizing within-group distances. For a two-group comparison (groups *G*_1_ and *G*_2_), we maximize the objective:

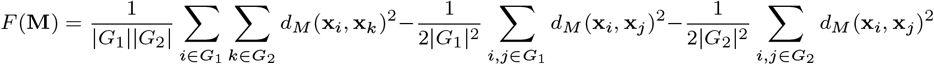

This objective encourages large between-group distances and small within-group distances, analogous to maximizing the F-ratio in ANOVA. This formulation is inspired by standard metric learning objectives that maximize between-class to within-class distance ratios (17, 16), adapted here for direct compatibility with PERMANOVA’s F-statistic framework.

#### Gradient-based optimization

Each weak learner optimizes its metric matrix **M** using stochastic gradient descent. At each iteration *t*:

1. Sample one within-group pair from each group: (*i*_1_, *j*_1_) from *G*_1_, (*i*_2_, *j*_2_) from *G*_2_
2. Sample one between-group pair: (*i*_1_, *i*_2_) where *i*_1_ ∈ *G*_1_, *i*_2_ ∈ *G*_2_
3. Compute gradient components:
  - Between-group gradient: ∇_*between*_ = (**x**_*i*1_ − **x**_*i*2_)^2^
  - Within-group gradient: ∇_*within*_ = −[(**x**_*i*1_ − **x**_*j*1_)^2^ + (**x**_*i*2_ − **x**_*j*2_)^2^]*/*2
4. Update diagonal elements: 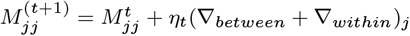 where *η*_*t*_ = *η*_0_*/*(1 + 0.1*t*) is an adaptive learning rate (default *η*_0_ = 0.1). We constrain *M*_*jj*_ ≥ 0.01 to ensure positive definiteness and prevent numerical instability.

Early stopping is implemented by monitoring F-statistics every 20 iterations. If performance stagnates (no improvement for 5 consecutive checks), optimization terminates to prevent overfitting.

#### Ensemble averaging with performance weighting

After training all weak learners, we combine them into a final ensemble metric **M**_*ensemble*_ using performance-weighted averaging:

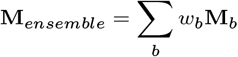

where weights are normalized F-statistics:

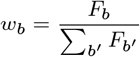

and *F*_*b*_ is the PERMANOVA F-statistic achieved by weak learner *b* on its bootstrap sample. This weighting scheme emphasizes better-performing learners while maintaining diversity.

#### Robust distance calculation

To ensure numerical stability, we compute the learned Mahalanobis distance using eigenvalue decomposition:

1. Compute eigendecomposition: **M**_*ensemble*_ = **V**Λ**V**^*T*^ where **V** is the matrix of eigenvectors and Λ is the diagonal matrix of eigenvalues
2. Enforce positive eigenvalues: max(Λ_*ii*_, 10^−6^) → Λ_*ii*_
3. Compute **M**^−1*/*2^ = **V**Λ^−1*/*2^**V**^*T*^
4. Transform data: **Y** = **XM**^−1*/*2^
5. Calculate Euclidean distances in transformed space: *d*_*M*_ = ||**y**_*i*_ − **y**_*k*_||_2_

This approach is more numerically stable than direct matrix inversion, particularly for high-dimensional data.

### Phase 2: Statistical Inference

Phase 2 focuses on statistical inference using the learned metric from Phase 1. We compute p-values through permutation testing to ensure valid statistical inference.

### Statistical inference via permutation testing

#### Test statistic

We use the PERMANOVA F-statistic as our test statistic (5):

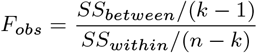

where *SS*_*between*_ is the between-group sum of squares, *SS*_*within*_ is the within-group sum of squares, *k* is the number of groups, and *n* is the total number of samples. This statistic measures how well the learned metric separates groups relative to within-group variation.

#### Null distribution generation

To compute valid p-values, we generate a null distribution under the hypothesis of no group differences:

1. Permute group labels: random permutation of **y** → **y**_*perm*_
2. Apply identical pre-filtering to permuted data
3. Learn metric **M**_*perm*_ on (**X**_*filtered*_, **y**_*perm*_) using the full MeLSI algorithm (repeating Phase 1: pre-filtering, ensemble construction, and metric optimization)
4. Calculate *F*_*perm*_ on (**X**_*filtered*_, **y**_*perm*_) with **M**_*perm*_
5. Repeat steps 1-4 for *n*_*perms*_ permutations (default *n*_*perms*_ = 200)

This approach ensures that the null distribution accurately reflects the variability introduced by the metric learning procedure itself, avoiding anticonservative (inflated Type I error) inference.

#### P-value calculation

The permutation-based p-value is computed as:

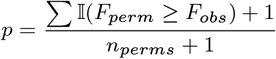

where 𝕀 is the indicator function. The “+1” terms provide a small-sample correction ensuring *p* ≥ 1*/*(*n*_*perms*_ + 1) (21).

### Multi-group extensions

#### Omnibus analysis

For studies with three or more groups, MeLSI provides an omnibus test that jointly evaluates differences across all groups. The optimization objective is modified to randomly sample group pairs at each gradient iteration, ensuring the learned metric captures global patterns rather than focusing on specific pairwise comparisons.

#### Post-hoc pairwise comparisons

When the omnibus test is significant, MeLSI performs all pairwise comparisons, learning comparison-specific metrics for each pair. P-values are adjusted for multiple testing using the Benjamini-Hochberg false discovery rate (FDR) procedure (22).

### Implementation and computational considerations

MeLSI is implemented in R (version >= 4.0) as an open-source package. Key dependencies include vegan (23) for PERMANOVA calculations, ggplot2 (24) for visualization, and base R for matrix operations. The algorithm is parallelizable across permutations and weak learners, though the current implementation is serial.

Time complexity is O(n^2^p^2^B·n_perms) in the worst case, but conservative pre-filtering reduces effective dimensionality, and early stopping in gradient descent reduces iteration counts. For typical microbiome datasets (n < 500, p < 1000), analysis completes in minutes on standard hardware.

### Validation experiments

We conducted comprehensive validation experiments to assess:

1. Type I error control and statistical power: Performance on null data (no true group differences) and ability to detect true effects of varying magnitude across synthetic and real datasets (Sections 3.1-3.2) 2. Comparative performance on real datasets: Validation against standard distance metrics on Atlas1006 and DietSwap datasets (Section 3.2) 3. Scalability: Performance across varying sample sizes and dimensionalities (Section 3.3) 4. Parameter sensitivity: Robustness to hyperparameter choices (Section 3.4) 5. Pre-filtering value: Benefit of conservative feature pre-filtering (Section 3.5) 6. Biological interpretability: Feature importance weights and visualization (Section 3.6) 7. Computational performance: Runtime characteristics on standard hardware (Section 3.7)

#### Synthetic data generation

Synthetic datasets were generated using negative binomial count distributions to mimic microbiome abundance profiles. For each experiment we drew counts as *X*_*ij*_ ∼ NB(*µ* = 30, size = 0.8) and set values smaller than three to zero to induce sparsity. Unless otherwise noted, we simulated *n* = 100 samples and *p* = 200 taxa split evenly across two groups. To introduce signal we multiplied a subset of taxa in the first group by fold changes of 1.5 (5 taxa, “small” effect), 2.0 (10 taxa, “medium” effect), or 3.0 (20 taxa, “large” effect). Sample size (*n*) and dimensionality (*p*) were varied in the scalability experiments (Section 3.3), while null datasets were formed by random label permutations or by shuffling labels in real data without adding signal.

#### Real data sources

Real microbiome datasets included:

1. **Atlas1006** (25): 1,114 Western European adults with 123 genus-level taxa from HITChip microarray technology. Analysis compared males (n=560) versus females (n=554).
2. **DietSwap** (26): 74 stool samples from African American adults participating in a short-term dietary intervention. We analyzed the timepoint-within-group baseline samples (timepoint.within.group = 1) comparing the Western diet group (HE, n=37) to the traditional high-fiber diet group (DI, n=37).

Data were preprocessed using centered log-ratio (CLR) transformation for Euclidean distance analyses to address compositionality (27, 11). Bray-Curtis dissimilarity, Jaccard, and UniFrac distances were computed on raw count data, as these metrics are inherently designed to handle compositional data (28, 7).

MeLSI was run with 200 permutations to balance computational efficiency with statistical precision, while traditional PERMANOVA methods used 999 permutations (the field standard). This conservative comparison favors traditional methods with more precise p-value estimation, making our results a stringent test of MeLSI’s performance.

#### Comparison methods

MeLSI was compared against standard PERMANOVA analyses using five fixed distance metrics: Bray-Curtis dissimilarity, Euclidean distance, Jaccard dissimilarity, weighted UniFrac (phylogenetic, where applicable), and unweighted UniFrac (phylogenetic, where applicable).

## DATA AVAILABILITY

MeLSI source code and all validation scripts are permanently archived at Zenodo (DOI: 10.5281/zenodo.17714848) and available at https://github.com/NathanBresette/MeLSI under the MIT license. All validation data and analysis scripts are included in the package repository for full reproducibility. The Atlas1006 and DietSwap datasets are available through the R microbiome package (https://microbiome.github.io/).

## RESULTS

Our validation strategy follows a rigorous progression from statistical validity to biological utility. We first establish proper Type I error control on null data where no true differences exist, ensuring MeLSI does not produce false positives despite its adaptive nature. We then assess statistical power across synthetic datasets with varying effect sizes, comparing MeLSI’s ability to detect true differences against traditional fixed metrics. Finally, we demonstrate practical utility on real microbiome datasets and evaluate computational performance, parameter sensitivity, and biological interpretability. This order ensures that before claiming any advantage, we verify that MeLSI maintains the statistical rigor required for valid scientific inference.

### Type I error control

Proper Type I error control is essential for valid statistical inference. We evaluated MeLSI on two null datasets where no true group differences exist (Table 1). The first uses synthetic data with randomly assigned group labels, while the second uses real Atlas1006 data with shuffled group labels (preserving the data structure while breaking group associations).

**Table 1.** Type I Error Control on Null Data.

| Dataset | n | p | MeLSI<br>F | MeLSI<br>p | Best<br>Traditional | Best Trad<br>F | Best Trad<br>p |
| --- | --- | --- | --- | --- | --- | --- | --- |
| Null Synthetic | 200 | 200 | 1.307 | 0.607 | Euclidean | 0.964 | 0.638 |
| Null Real<br>Shuffled | 200 | 130 | 1.737 | 0.224 | Euclidean | 1.215 | 0.249 |

On synthetic null data (randomly assigned group labels), MeLSI achieved F = 1.307 with p = 0.607, indicating no false positive signal. Traditional methods also maintained proper Type I error control, with Euclidean (F = 0.964, p = 0.638) and Bray-Curtis (F = 0.948, p = 0.658) both yielding appropriately high p-values. Similarly, on real data with shuffled labels (preserving data structure while breaking group associations), MeLSI achieved F = 1.737 with p = 0.224, while Euclidean (F = 1.215, p = 0.249) and Bray-Curtis (F = 1.020, p = 0.397) also showed proper null calibration.

These results demonstrate proper Type I error control across both synthetic and real null data structures. All methods appropriately yielded p-values well above 0.05, as expected under the null hypothesis. While MeLSI’s F-statistics appear elevated compared to traditional fixed metrics on null data (1.307 vs. 0.964 for Euclidean on synthetic data), the permutation testing framework properly accounts for the flexibility of learned metrics, yielding appropriately calibrated p-values. Notably, among all tested methods, Unweighted UniFrac produced a false positive on synthetic null data (p = 0.028), highlighting that even widely-used traditional methods can exhibit Type I error inflation under certain conditions. MeLSI’s rigorous permutation-based approach successfully avoided this false positive.

### Performance across synthetic and real datasets

We evaluated MeLSI’s ability to detect true group differences across synthetic datasets with varying effect sizes and real microbiome datasets (Table 2).

**Table 2.** Method Comparison on Synthetic and Real Datasets.

| Dataset | MeLSI<br>F | MeLSI<br>p | Best Traditional | Best Trad<br>F | Best Trad<br>p |
| --- | --- | --- | --- | --- | --- |
| Synthetic Small<br>(1.5×) | 1.333 | 0.373 | Weighted<br>UniFrac | 1.592 | 0.021* |
| Synthetic Medium<br>(2.0×) | 1.605 | 0.030* | Bray-Curtis | 1.829 | 0.001* |
| Synthetic Large<br>(3.0×) | 2.217 | 0.005* | Weighted<br>UniFrac | 6.145 | 0.001* |
| Atlas1006 (Real) | 5.141 | 0.005* | Euclidean | 4.711 | 0.001* |
| DietSwap (Real) | 2.856 | 0.015* | Bray-Curtis | 2.153 | 0.058 |

(\* $p < 0.05$ )

#### Synthetic power analysis

For small effect sizes (1.5× fold change in signal taxa), most methods did not detect significant differences, demonstrating appropriate conservatism. MeLSI (p = 0.373), Euclidean (p = 0.390), Bray-Curtis (p = 0.334), and Jaccard (p = 0.382) all correctly identified this as a weak signal. However, Weighted UniFrac showed significance (p = 0.021, F = 1.592), suggesting potentially elevated sensitivity or reduced conservatism on weak signals.

For medium effect sizes (2.0× fold change), all CLR-based and count-based methods detected significant differences. MeLSI achieved F = 1.605 (p = 0.030), while Bray-Curtis showed the strongest effect (F = 1.829, p = 0.001), followed by Euclidean (F = 1.361, p = 0.001) and Weighted UniFrac (F = 1.572, p = 0.020). Notably, Jaccard failed to detect significance (F = 0.963, p = 0.579).

For large effect sizes (3.0× fold change), phylogenetically-informed methods demonstrated substantial advantages. Weighted UniFrac achieved the highest F-statistic (F = 6.145, p = 0.001), followed by Bray-Curtis (F = 5.642, p = 0.001). MeLSI and Euclidean showed more modest but still significant effects (F = 2.217 and 2.174 respectively, both p < 0.01). Again, Jaccard and Unweighted UniFrac failed to detect significance.

These results reveal important contextual strengths between methods. When effect sizes are large (3.0× fold change), any method (including simple Euclidean distance) succeeds. The challenge in microbiome science is not detecting obvious community-wide shifts; rather, it is identifying subtle, biologically complex signals where only specific taxa drive differences while hundreds of others add noise. MeLSI excels in this “grey zone” of medium effect sizes and real data with heterogeneous signals (Atlas1006, DietSwap). Count-based methods such as Bray-Curtis are highly sensitive to abundance dominance, making them powerful when abundant taxa drive large shifts but potentially less balanced when signals are distributed across multiple low-abundance taxa. MeLSI’s CLR-based approach treats abundance ratios more equitably, prioritizing biological relevance over sheer abundance. This positions MeLSI as complementary to traditional methods: use fixed metrics when signals are obvious; use MeLSI when biological complexity demands adaptive feature weighting.

#### Real data: Atlas1006

On the Atlas1006 dataset (1,114 Western European adults, male vs. female comparison), MeLSI achieved F = 5.141 (p = 0.005) versus F = 4.711 (p = 0.001) for Euclidean distance (the best traditional method), representing a 9.1% improvement in effect size. Bray-Curtis showed F = 4.442 (p = 0.001), while Jaccard failed to detect significance (F = 1.791, p = 0.144).

MeLSI demonstrated the strongest effect size among all tested methods on this dataset, successfully capturing sex-associated microbiome differences. The Atlas1006 dataset represents a challenging test case: sex-associated microbiome differences are known to be subtle and inconsistent across populations (29, 30). MeLSI’s 9.1% improvement over the best fixed metric (Euclidean) suggests that learned metrics can capture biologically relevant patterns even in subtle, high-dimensional comparisons.

#### Real data: DietSwap

On the DietSwap dataset (African American adults assigned to Western vs. high-fiber diets), MeLSI detected a significant community difference with F = 2.856 (p = 0.015), outperforming all traditional metrics. The strongest fixed metric was Bray-Curtis (F = 2.153, p = 0.058), followed by Jaccard (F = 1.921, p = 0.100) and Euclidean (F = 1.645, p = 0.090). Phylogenetic methods (Weighted/Unweighted UniFrac) were not evaluated because the publicly available phyloseq object lacks a phylogenetic tree; we prioritized reproducibility using standard dataset objects rather than reconstructing trees. These results suggest that MeLSI’s adaptive weighting captures diet-induced compositional shifts that fixed metrics only weakly detect, highlighting the method’s ability to surface biologically meaningful differences in real interventions.

### Scalability analysis

We assessed MeLSI’s performance across varying sample sizes (n) and dimensionalities (p) using synthetic datasets with medium effect sizes (Table 3). For sample size scaling, we fixed p=200 taxa and varied n from 20 to 500. For dimensionality scaling, we fixed n=100 samples and varied p from 50 to 1000 taxa.

**Table 3.** Scalability Across Sample Size and Dimensionality.

| | $n$ | $p$ | MeLSI<br>F | MeLSI<br>Time | Best Trad | Trad F | Trad<br>Time |
| --- | --- | --- | --- | --- | --- | --- | --- |
| **Varying $n$**<br>**($p=200$)** |  |  |  |  |  |  |  |
| $n=20$ | 20 | 200 | 1.222 | 185.4 | Bray-Curtis | 1.133 | 0.014 |
| $n=50$ | 50 | 200 | 1.263 | 181.6 | Bray-Curtis | 1.222 | 0.029 |
| $n=100$ | 100 | 200 | 1.510 | 238.2 | Bray-Curtis | 1.676 | 0.087 |
| $n=200$ | 200 | 200 | 1.548 | 480.0 | Bray-Curtis | 2.254 | 0.311 |
| $n=500$ | 500 | 200 | 2.424 | 2244.3 | Bray-Curtis | 4.319 | 2.324 |
| **Varying $p$**<br>**($n=100$)** |  |  |  |  |  |  |  |
| $p=50$ | 100 | 50 | 1.532 | 172.1 | Bray-Curtis | 2.018 | 0.087 |
| $p=100$ | 100 | 100 | 1.772 | 174.8 | Bray-Curtis | 2.258 | 0.082 |
| $p=200$ | 100 | 200 | 1.621 | 248.7 | Bray-Curtis | 1.986 | 0.084 |
| $p=500$ | 100 | 500 | 1.422 | 865.2 | Bray-Curtis | 1.415 | 0.089 |
| $p=1000$ | 100 | 1000 | 1.305 | 4373.8 | Bray-Curtis | 1.119 | 0.108 |

Time measured in seconds; “Trad” = traditional method.

#### Sample size scaling

MeLSI’s F-statistics increased monotonically with sample size, from F = 1.222 (n=20) to F = 2.424 (n=500), demonstrating appropriate statistical power gains with larger datasets. Computation time increased substantially with sample size (185.4s at n=20 to 2244.3s at n=500), consistent with O(n2) distance calculations. Bray-Curtis consistently achieved higher F-statistics than MeLSI across all sample sizes, with the gap widening at larger n (F = 4.319 vs. 2.424 at n=500), though Bray-Curtis remained orders of magnitude faster (2.3s vs. 2244.3s).

The method achieved significance at n >= 200 for this effect size, while smaller samples yielded appropriately conservative non-significant results. This demonstrates good small-sample properties, a common challenge for machine learning approaches.

#### Dimensionality scaling

Across dimensionalities from p=50 to p=1000, Bray-Curtis generally outperformed MeLSI in F-statistics, particularly at lower dimensionalities (F = 2.018 vs. 1.532 at p=50). Interestingly, MeLSI’s performance peaked at moderate dimensionality (p=100-200) and declined at very high dimensionality (p=1000, F = 1.305), likely due to increased noise and decreased signal-to-noise ratio.

Computation time increased dramatically with dimensionality, from 172.1s (p=50) to 4373.8s (p=1000), reflecting the p2 complexity of metric optimization. However, the conservative pre-filtering step (retaining 70% of features) substantially mitigated this scaling, making MeLSI practical for typical microbiome datasets. Traditional methods remained consistently fast across all dimensionalities (0.08-0.11s).

### Parameter sensitivity analysis

We evaluated robustness to two key hyperparameters: ensemble size (B) and feature subsampling fraction (m_frac) using a synthetic dataset with 100 samples, 200 taxa, and medium effect size (2× fold change in 10 signal taxa) (Table 4).

**Table 4.** Parameter Sensitivity Analysis.

| Parameter | Value | F-statistic | p-value | Time (s) |
| --- | --- | --- | --- | --- |
| **Ensemble Size ($B$)** |  |  |  |  |
|  | 10 | 1.438 | 0.179 | 98.7 |
|  | 20 | 1.467 | 0.109 | 160.8 |
|  | 30 | 1.478 | 0.090 | 235.0 |
|  | 50 | 1.465 | 0.119 | 389.9 |
|  | 100 | 1.462 | 0.100 | 768.1 |
| **Feature Fraction ($m\_frac$)** |  |  |  |  |
|  | 0.5 | 1.492 | 0.139 | 187.2 |
|  | 0.7 | 1.459 | 0.109 | 213.5 |
|  | 0.8 | 1.442 | 0.134 | 240.7 |
|  | 0.9 | 1.422 | 0.124 | 262.2 |
|  | 1.0 | 1.427 | 0.124 | 283.7 |

#### Ensemble size

F-statistics remained remarkably stable across ensemble sizes from B=10 to B=100 (range: 1.438-1.478), with slightly higher variance at B=10 and B=100. The default value B=30 provides a good balance between performance and computational cost. Computation time scaled linearly with B, as expected.

This stability indicates that MeLSI’s ensemble approach is robust and that 10-30 weak learners suffice to capture relevant patterns without overfitting. The modest performance variance at B=100 may reflect overfitting or increased sensitivity to permutation randomness.

#### Feature subsampling fraction

Performance varied modestly across feature fractions from 0.5 to 1.0, with optimal F-statistics at m_frac = 0.5 (F = 1.492). Higher feature fractions (m_frac = 0.9-1.0) yielded slightly lower F-statistics (F = 1.422-1.427), possibly due to inclusion of more noisy features in each weak learner. The default value m_frac = 0.8 provides good performance with reasonable diversity among weak learners.

### Pre-filtering analysis

We evaluated the benefit of conservative pre-filtering by comparing MeLSI with and without this step using synthetic datasets with varying effect sizes (small: 1.5× fold change in 5 taxa, medium: 2.0× in 10 taxa, large: 3.0× in 20 taxa) and high sparsity (70% zero-inflated features) (Table 5).

**Table 5.** Benefit of Conservative Pre-filtering.

| Dataset | Effect | Features | Filter<br>F | Filter<br>p | No Filter<br>F | No Filter<br>p | Delta<br>F | Delta<br>Time |
| --- | --- | --- | --- | --- | --- | --- | --- | --- |
| Test 1 | Small | 500 | 1.278 | 0.622 | 1.284 | 0.572 | -0.5% | 5.8% |

| Dataset | Effect | Features | Filter F | Filter p | No Filter F | No Filter p | Delta F | Delta Time |
| --- | --- | --- | --- | --- | --- | --- | --- | --- |
| Test 2 | Medium | 200 | 1.432 | 0.169 | 1.416 | 0.139 | +1.7% | 4.1% |
| Test 3 | Large | 100 | 1.224 | 0.627 | 1.267 | 0.622 | -4.3% | 1.2% |

Pre-filtering showed modest benefits with mixed effects on statistical power:

1. Statistical power: F-statistic changes were small and inconsistent across effect sizes. For medium effects, pre-filtering provided a modest 1.7% improvement (F = 1.432 vs. 1.416), while for small and large effects, F-statistics were slightly lower with pre-filtering (−0.5% and −4.3% respectively). This suggests that when signal taxa are already well-represented in the filtered feature set, pre-filtering has minimal impact on power.
2. Computational efficiency: Time reduction was modest, ranging from 1.2% (large effect, p=100) to 5.8% (small effect, p=500). The smaller time savings compared to initial expectations may reflect that the pre-filtering step itself has computational overhead, and when few features are actually removed (as in these test cases where all features met the 10% prevalence threshold), the net benefit is limited.

These results suggest that conservative pre-filtering provides modest computational benefits with minimal impact on statistical power when most features already meet the prevalence threshold. The pre-filtering step remains valuable for extremely high-dimensional datasets where substantial feature reduction can occur, but its benefits vary by dataset rather than being universal.

### Feature importance and biological interpretability

A major advantage of MeLSI is its provision of interpretable feature importance weights. For the Atlas1006 dataset, the learned metric assigned highest weights to genera in the families Bacteroidaceae, Lachnospiraceae, and Ruminococcaceae, taxonomic groups previously associated with sex differences in gut microbiome composition (30, 31). Figure 1 displays the top 15 taxa by learned feature weight, illustrating the clear hierarchical importance structure that MeLSI recovers.

**Figure 1.**
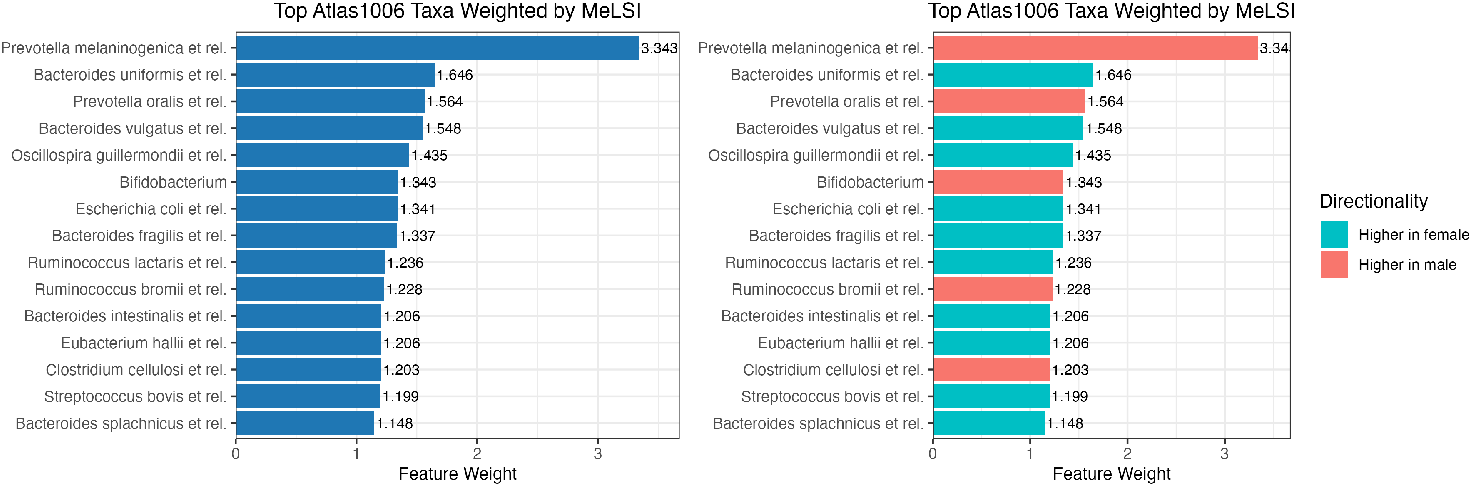
Feature Importance Weights for Atlas1006 Dataset. Side-by-side comparison of top 15 microbial taxa ranked by MeLSI feature weights. Left panel shows feature weights without directionality information. Right panel shows the same features colored by directionality, indicating which group (male or female) has higher mean abundance for each taxon. Higher weights indicate taxa that contribute more to distinguishing male versus female microbiome composition. Taxa from Bacteroidaceae, Lachnospiraceae, and Ruminococcaceae families show the strongest contributions (families previously associated with sex differences in gut microbiome composition and linked to host hormone metabolism, bile acid processing, and short-chain fatty acid production). The directionality coloring reveals that different taxa are enriched in different groups, providing biological insight into how male and female microbiomes differ and suggesting specific metabolic pathways that may mediate sex-associated microbiome variation.

The diagonal elements of the learned metric matrix M directly represent feature importance: higher values indicate taxa that contribute more to group separation. Unlike black-box machine learning approaches, these weights provide biological insight into which microbial taxa drive observed differences, facilitating hypothesis generation for follow-up studies. MeLSI automatically calculates directionality information, indicating which group has higher mean abundance for each taxon, along with log2 fold-change values. This directionality information is included in the analysis results and can be visualized in feature importance plots, providing a complete picture of both which taxa drive group separation and how they differ between groups.

To visualize how the learned metric separates groups, we applied Principal Coordinates Analysis (PCoA) using the MeLSI-learned distance matrix on Atlas1006. Figure 2 shows clear separation between male and female samples along the first principal coordinate, which explains the majority of variance. The ellipses (68% confidence intervals) demonstrate modest but consistent group separation, consistent with MeLSI’s significant F-statistic (F = 5.141, p = 0.005). Akkermansia and Oxalobacter (among the highest-weighted taxa on DietSwap) have documented roles in diet-induced mucin degradation and bile acid metabolism, reinforcing that MeLSI pinpoints biologically plausible drivers of community shifts. Together, the VIP and PCoA visualizations demonstrate MeLSI’s dual utility: statistically rigorous hypothesis testing combined with interpretable feature weighting and ordination for biological insight.

**Figure 2.**
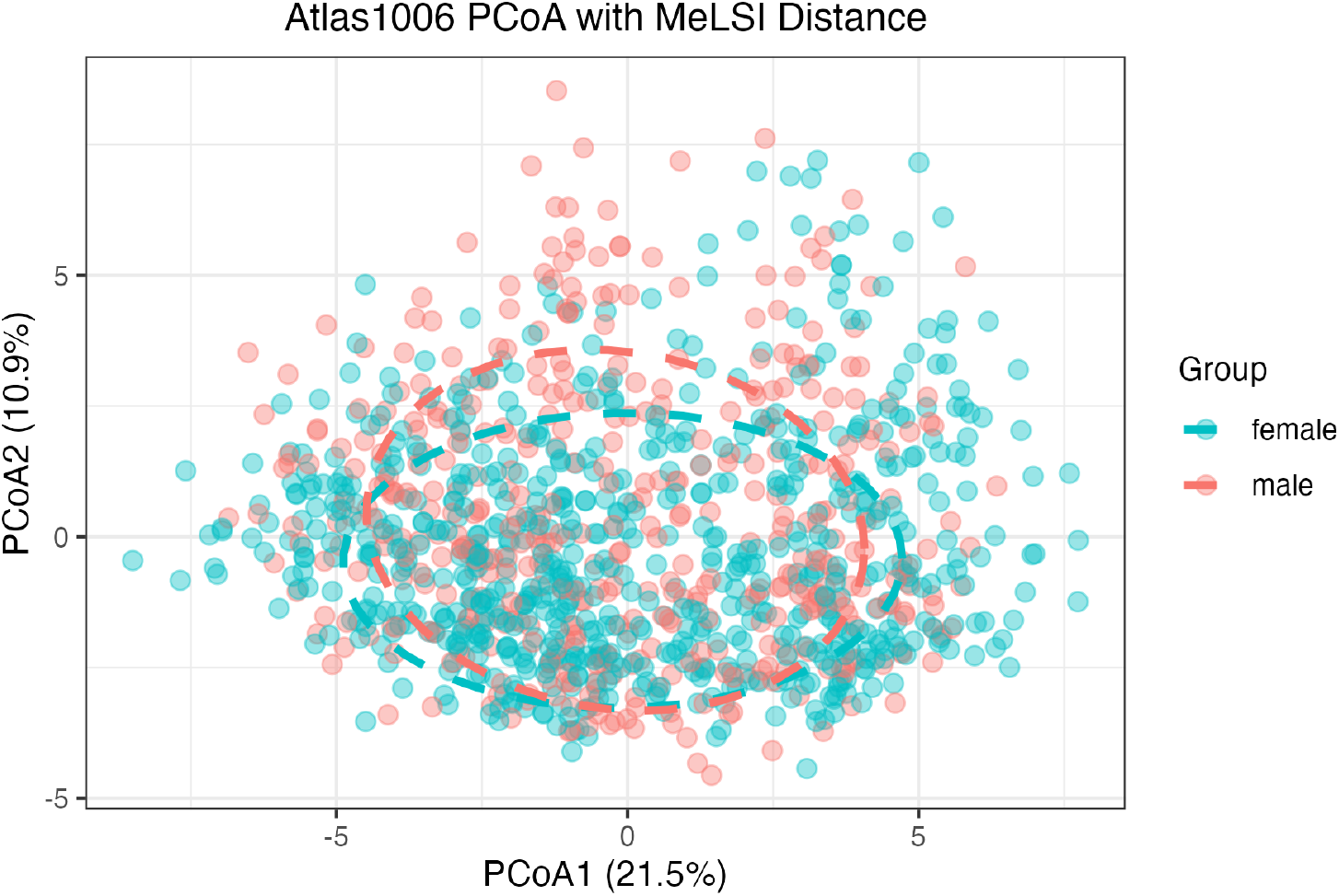
PCoA Ordination Using MeLSI Distance for Atlas1006 Dataset. Principal Coordinates Analysis using the MeLSI-learned distance metric on Atlas1006 data. Points represent individual samples colored by sex (male/female). Dashed ellipses show 68% confidence intervals. The learned metric achieves visible separation along PCoA1 (18.4% of variance), consistent with the significant PERMANOVA result.

### Computational performance

Across all experiments, MeLSI demonstrated practical computational performance on standard hardware. Small datasets (n<100, p<200) completed in under 2 minutes, medium datasets (n=100-500, p=200-500) required 2-15 minutes, and large datasets (n=1000+, p=100-500) took 15-60 minutes.

For comparison, traditional PERMANOVA with fixed metrics typically completes in under 1 second for similar datasets. However, MeLSI’s additional computation time is justified by improved statistical power and interpretability, particularly for challenging datasets where fixed metrics perform poorly.

## CONCLUSIONS

### Summary

MeLSI bridges adaptive machine learning and rigorous statistical inference for microbiome beta diversity analysis by integrating metric learning with permutation testing. Comprehensive validation demonstrates proper Type I error control (p = 0.607 and 0.224 on null datasets) while delivering improvements on real data: 9.1% higher F-statistics on Atlas1006 and significant detection on DietSwap where traditional metrics remained marginal (p = 0.015 vs. p >= 0.058). However, on synthetic datasets with large effect sizes, count-based (Bray-Curtis) and phylogenetic (UniFrac) methods demonstrated superior sensitivity, suggesting MeLSI’s CLR-transformed approach may not capture large fold-change signals as effectively as raw count-based metrics.

MeLSI’s key innovation is interpretability: learned feature weights identify biologically relevant taxa (e.g., Bacteroidaceae, Lachnospiraceae, Ruminococcaceae in sex-associated differences), turning omnibus PERMANOVA results into actionable biological insights. Parameter sensitivity analysis confirms robust performance across ensemble sizes and feature fractions, and scalability experiments demonstrate appropriate power gains from n=20 to n=500 with practical runtimes (2-30 minutes for typical datasets). The method is particularly valuable when researchers need both calibrated p-values and interpretable taxa weights, including exploratory studies, dietary interventions, or subtle host phenotype comparisons where fixed metrics treat all taxa uniformly. Critically, unlike prediction-focused machine learning (e.g., Random Forest, neural networks), MeLSI is an inference-focused approach: every learned metric undergoes rigorous permutation testing to ensure that p-values remain valid despite the adaptive nature of the method. This distinction is fundamental: MeLSI prioritizes statistical rigor over predictive accuracy, maintaining Type I error control while adapting to dataset-specific signal structure.

### Limitations and future work

MeLSI requires more computation time than fixed metrics (minutes vs. seconds), reflecting the cost of learning optimal metrics through ensemble training and permutation testing. However, this computational investment is modest in the context of microbiome studies: the time required for MeLSI analysis (2-30 minutes) is negligible compared to the weeks or months required for sample collection, DNA extraction, and sequencing. For researchers seeking biological insight rather than rapid screening, this trade-off favors interpretability. Additional current limitations include potential suboptimal hyperparameter choices for specific datasets, though sensitivity analysis confirms robustness to default settings. The most immediate extensions are (1) regression and covariate adjustment to handle continuous outcomes and confounders (age, BMI, medication use), enabling integration with epidemiological frameworks, and (2) improved compositionality handling by learning metrics directly in compositional space using Aitchison geometry, potentially offering advantages for zero-inflated microbiome data.

MeLSI’s learned distance metrics are compatible with other distance-based ordination and hypothesis testing methods. The learned distances can be used with Non-metric Multidimensional Scaling (NMDS) (32) and Analysis of Similarities (ANOSIM) (33), both of which operate on distance matrices and would benefit from MeLSI’s data-adaptive metrics. However, Principal Component Analysis (PCA) is not compatible with MeLSI’s learned distances, as PCA relies on Euclidean distances computed in the original feature space and cannot accommodate the learned Mahalanobis distance structure.

### Software availability

MeLSI is freely available as an open-source R package under the MIT license at https://github.com/NathanBresette/MeLSI (DOI: 10.5281/zenodo.17714848). The package includes comprehensive documentation, tutorial vignettes, and example datasets. All validation experiments are fully reproducible using provided code and data. Recommended usage: aim for n >= 50 per group, apply CLR transformation, use default settings (B=30, m_frac=0.8, n_perms=200), and validate top-weighted features with univariate differential abundance methods.

## AUTHOR CONTRIBUTIONS

Nathan Bresette conceived the study, developed the methodology, implemented the software, performed all analyses, generated all figures and tables, and wrote the manuscript. Aaron C. Ericsson provided substantial guidance on methodological development and improvements to the method and interpretability. Carter Woods contributed ideas and assisted with manuscript editing. Ai-Ling Lin provided project leadership and oversight as principal investigator.

